# Aneuploidy promotes intestinal dysplasia in Drosophila

**DOI:** 10.1101/280768

**Authors:** Luís Pedro Resende, Augusta Monteiro, Rita Brás, Tatiana Lopes, Claudio E. Sunkel

## Abstract

Aneuploidy is associated with different human diseases, particularly cancer, but how different cell types within tissues respond to aneuploidy is not fully understood. In some studies, aneuploidy has been shown to have a deleterious effect and lead to cell death, however it has also been shown to be a causal event of tumorigenesis in other contexts.

Here, we show that *Drosophila* intestinal stem cells have a particular tolerance to aneuploidy and do not activate apoptosis in response to chromosome misegregation like other non-stem cells. Instead, we observe the development of tissue dysplasia characterized by an accumulation of progenitor cells, increased stem cell proliferation rate, and an excess of cells of the enteroendocrine lineage. Our findings highlight the importance of mechanisms acting to prevent aneuploidy within tissue stem cells and provide an *in vivo* model of how these cells can act as reservoirs for genomic alterations that can lead to dysplasia.

## Introduction

Aneuploidy is characterized by the presence of an abnormal number of chromosomes in a cell and is a hallmark of different human diseases - it is one of the major causes of spontaneous miscarriages, a hallmark of cancer, and it has been linked to neurodegeneration and aging [1-2]. Aneuploidy is present in more than 90% of human tumors, but several studies report a detrimental effect of aneuploidy on cells leading to cell death or cell cycle arrest. [3]. This complexity is partially explained by the fact that the effects of aneuploidy seem to be cell type specific.

Tissue stem cells are responsible for the constant self-renewal of our tissues, and their behavior must be tightly regulated to prevent diseases. Contrasting with other proliferative non-stem cells [4-6], adult stem cells have been proposed to tolerate aneuploidy and not activate apoptosis in response to genomic instability [7-10]. This resistance to aneuploidy underscores a need to understand how tolerated aneuploidy impacts adult stem cell behavior and tissue homeostasis. Here, we show that *Drosophila* intestinal stem cells (ISCs) are competent for the Spindle-Assembly Checkpoint (SAC), a surveillance mechanism that ensures faithful chromosome segregation during mitosis [11]. However, while we find that prolonged SAC impairment results in induction of aneuploidy in intestinal progenitor cells, this does not lead to apoptosis activation and, consequently, loss of these cells. Instead, we observe tissue dysplasia characterized by an accumulation of progenitor cells, increased stem cell proliferation rate, and an excess of cells of the enteroendocrine lineage. Importantly, these phenotypes are recapitulated when aneuploidy is induced via manipulation of other biological mechanisms associated with aneuploidy, like defects on kinetochore structure or centrosome amplification, suggesting they portrait a broad effect of aneuploidy on ISCs. Our findings describe an *in vivo* model on how a failure to maintain a correct genomic content can lead to stem cell malfunction and tissue pathology.

## Results and discussion

### *Drosophila* intestinal stem cells are SAC competent

The *Drosophila* intestine is a powerful model system to study adult stem behavior *in vivo*, where markers are available for all cell types that compose the intestinal epithelium (Figure 1 A), and a diversity of genetic tools can be used to manipulate gene expression in a cell-type and temporally-controlled manner [12]. In the *Drosophila* posterior midgut, multipotent ISCs and enteroblasts (EBs) constitute the progenitor cell population of this tissue. Differentiated cell types in the adult midgut include secretory enteroendocrine (EE) cells, and absorptive polyploid enterocytes (EC). ISCs have the potential to divide symmetrically or asymmetrically with regards to cell fate. When dividing asymmetrically they can give rise to either an EB or an EE, a process regulated by bidirectional Notch signaling [13-17]. ECs are generated through differentiation of EBs [18-19].

**Figure 1.**
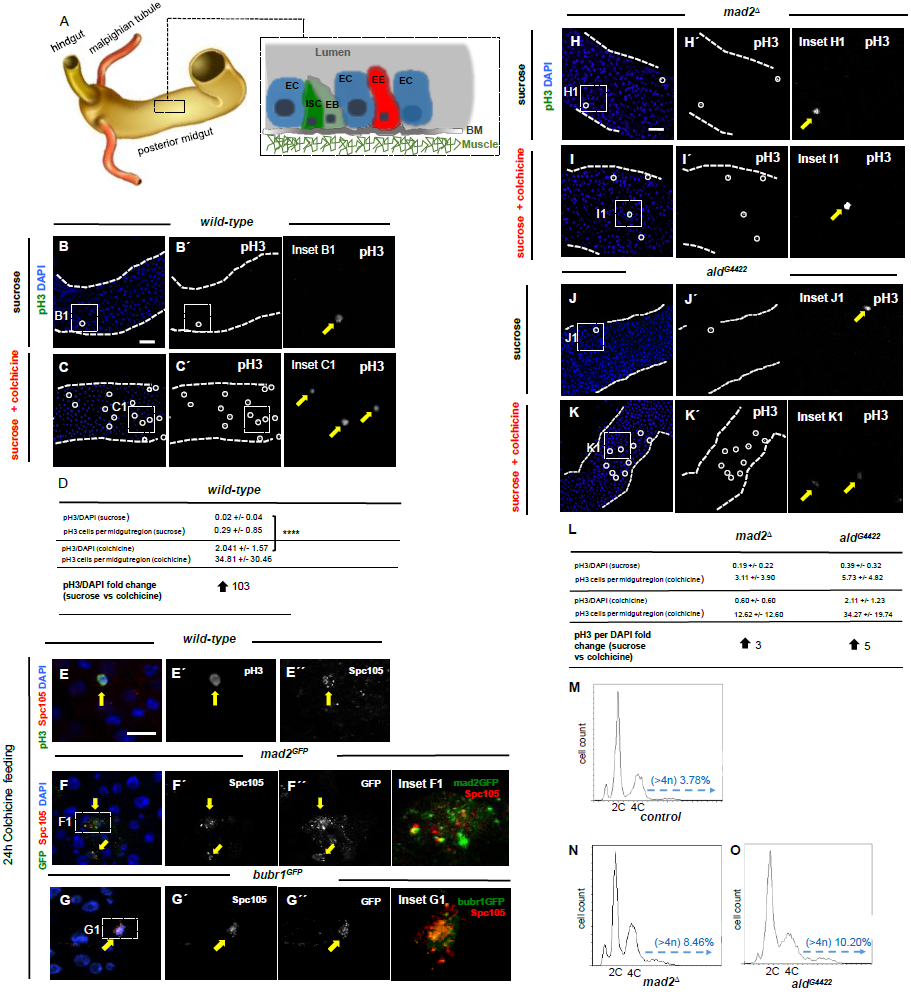
Drosophila Intestinal stem cells are competent for the spindle-assembly checkpoint. **A)** Anatomic organization of the adult *Drosophila* intestine and schematic representation of different cell types found in the posterior midgut (black dashed box); ISCs/EBs in green are the progenitor cells and are found in close association with basement membrane (BM); Differentiated cell types include Enteroendocrine cells (EE, red) and absorptive enterocytes (EC, blue); **B)** Mitotic cells labeled with phospo-histone 3 (pH3, Green, B´) in *wild-type* 2-5 day-old OreR fed with 5% sucrose control solution during 24h (white circle and yellow arrow show pH3 positive cell, inset B1); **C)** same as B) but fed with 5% sucrose and 0.2mg/mL colchicine during the same period; note clear increase in pH3 positive cells (compare C´ with B´); **D)** Quantification of pH3 positive cells present in the first two fields of view in the 40X objective of situations described in B) and C); **** p<0.0001, Mann–Whitney U test; N(intestines)=24 for sucrose, N(intestines)=47 for sucrose+colchicine; **E)** kinetochore marker Spc105 is detected in SAC arrested ISCs (pH3 positive, yellow arrow); **F) and G)** *mad2* or *bubr1* reporter lines show GFP signal in SAC arrested cells (yellow arrows); **H) to K)** 2-5 day-old *mad2* or *mps1* mutants flies fed with the same feeding method as described for *wild-type* flies in B) and C); white circles and yellow arrows show pH3 positive cells; **L)** Quantification of pH3 positive cells present in the first two fields of view in the 40X objective of situations described in H) and K); N(intestines)=22 for *mad2* mutants fed with sucrose, N(intestines)=29 for *mad2* mutants fed with sucrose+colchicine, N(intestines)=11 for *mps1* mutants fed with sucrose, N(intestines)=11 for Mps1 mutants fed with sucrose+colchicine; **M to O)** Examples of FACS profiles of GFP positive cells (ISCs/EBs) from control and mutants for *mad2* or *mps1*; for control and *mad2* mutants 5 biological replicates were performed and in all cases the percentage of aneuploidy was higher in the *mad2* mutants (p-value<0.01, Fisher exact t-test); for control and *mps1* mutants 3 biological replicates were performed, in all cases the percentage of aneuploidy was higher in the *mps1* mutants (p-value<0.01, Fisher exact t-test); average of % aneuploidy +/- st dev: controls 4.4 +/- 2.5, Mad2 mutants 8.9 +/- 3.7, Mps1 mutants 11.9 +/- 1.9; scale bars on B) = 40μm, panel C) and panels and H) to K) are on the same magnification; scale bars on E) = 4μm, panels F) and G) are on the same magnification.

In order to investigate if SAC impairment could be used as a strategy to induce aneuploidy in ISCs we first examined if SAC function was active in these cells. SAC competent cells respond to depolymerization of microtubules by drugs such as colchicine with a mitotic arrest in prometaphase [20]. Previous studies have reported that proliferative cells within the intestine arrest in mitosis when flies are fed with microtubule depolymerizing drugs like colchicine or colcemid [21-23]. To confirm that ISCs are able to respond to microtubule spindle depolymerization of microtubules we fed flies with colchicine. Consistent with ISCs being SAC functional, a high number of mitotic cells (phospho-histone H3 positive) was found in *wild-type* flies fed with colchicine for 24h, while these were rarely found in flies fed during the same period with a sucrose solution (Figure 1 B-D). In agreement, we observed that after colchicine feeding mitotic cells showed kinetochore accumulation of SAC proteins such as Mad2 and BubR1 (Figure 1 E-G) [24-25]. To further validate the SAC response in ISCs, we tested if SAC genes were essential for this prometaphase arrest. Unlike in mammals, *Drosophila* homozygous mutants for *mad2* SAC gene (*mad2*^*Δ*^) are viable and fertile [26]. Our laboratory characterized another SAC mutant (*ald*^*G4422*^) which encodes the *mps1* gene [27-28], that allows survival of a small percentage of homozygous flies until adult stages. Both of these SAC mutants are checkpoint defective and show high rates of aneuploidy in their neuroblast population [26-27]. To investigate if the SAC is impaired in these mutants, colchicine feeding experiments were performed on *mad2* and *mps1* homozygous mutant flies and the number of mitotic cells per total cells were compared to *wild-type* controls. In *wild-type* flies, colchicine feeding lead to a 103-fold increase on the number of mitotic cells per total cells, whereas in SAC mutants this increase was significantly lower, only of 3-fold for *mad2*^*Δ*^ and 5-fold for *ald*^*G4422*^ (Figure 1 H-L). Thus, we conclude that the colchicine arrest observed in *wild-type* ISCs is due to an active SAC. Furthermore, we could also observe that overexpression of the SAC gene *mps1* causes mitotic arrest consistent with its role in an active checkpoint response in ISCs (Figure S1 A-C).

In different model organisms and cell types, a defective checkpoint has consistently been associated with abnormal chromosome segregation and aneuploidy induction [11]. After showing that ISCs have a functional SAC response, we wanted to determine whether loss of the SAC could lead to aneuploidy induction in intestinal progenitor cells. To do so, we performed FACS analysis of cells isolated from the intestine of control or SAC mutant intestines labelled with the *esgGAL4,UASGFP* transgene that identifies both ISCs and EBs [18-19][29] (Figure 1 M-O). FACS analysis has been used successfully to detect increases in specific types of aneuploidy within *Drosophila* imaginal discs after SAC impairment [4-6] and, more recently, adult intestinal progenitor cells [30-31]. It is important to note that this method to detect aneuploidy is conservative and misses any aneuploidy classes with less than the equivalent of the G2 DNA content. As expected, homozygous mutant cells for *mad2* or *mps1* showed higher rates of aneuploidy within their GFP positive cell population (ISCs/EBs) with a DNA content >4n, confirming that loss of SAC genes results in aneuploidy induction in intestinal progenitor cells.

### SAC impairment results in intestinal dysplasia

To characterize the impact of aneuploidy induction in the stem cell population, we quantified the number of ISCs/EBs in control, *mad2* or *mps1* mutant flies. We observed a significant accumulation of ISCs/EBs in the intestines of *mad2* or *mps1* mutants when compared to controls (Figure 2 A-D). These observations are in sharp contrast to previous findings on the impact of aneuploidy in *Drosophila* imaginal cells, where aneuploidy induction results in apoptosis, and only if apoptosis is blocked, a tumorigenic phenotype can be observed [4-6]. Interestingly, it has been shown that aneuploidy does not induce embryonic stem cells death [32], and more recent studies have proposed that this resistance to aneuploidy might be conserved in different adult stem cell populations [7-9], including the *Drosophila* ISCs [10]. Recently, it has been proposed that aneuploidy has a detrimental effect on *Drosophila* brain and intestinal stem cell populations, leading to their loss through premature differentiation [33]. As experiments performed in the intestine by Gogendeu *et al* were based on a single genetic condition used to induce aneuploidy (*Bub3* loss of function), and the main effect of aneuploidy was different from our initial results with *mad2* and *mps1* mutants, we decided to analyze in greater detail the consequences of aneuploidy induction in intestinal adult stem cells using other genetic conditions.

**Figure 2.**
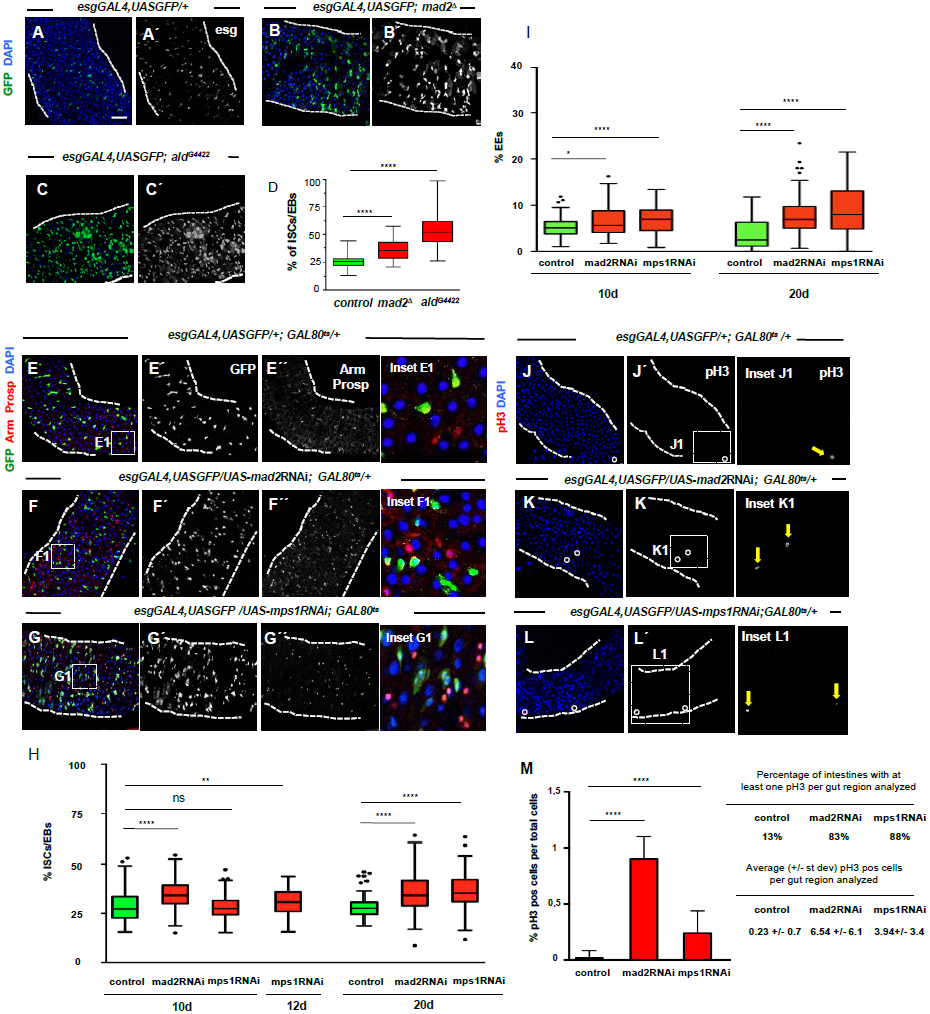
Mad2 and Mps1 loss-of-function results in intestinal dysplasia. **A)** 2-5 day-old control intestine where ISCs/EBs can be visualized by GFP expression under the control of the esgGAL4 driver, DAPI marks cell nuclei; **B) and C)** 2-5 day-old intestines of homozygous mutants for SAC genes *mad2* or *mps1*; note accumulation of ISCs/EBs (compare B´ or C´ with A´); **D)** Quantification of percentage of progenitor cells (ISCs/EBs, GFP positive) per total cell number in situations A) to C); **E)** 20 day-old control intestine where ISCs and EBs are marked based on *esg* expression (green, panel E´), EEs can be identified by expression of prospero (nuclear red signal, E´´), armadillo (Arm) marks all cells membrane, DAPI marks cell nuclei; **F) and G)** Intestines where RNAi constructs against SAC genes *mad2* or *mps1* were expressed under the control of the *esgGAL4* promoter during the first 20 days of the adult fly; note accumulation of ISCs/EBs (compare F´ and G´ with E’ or green cells in the corresponding insets); note also the excess of EEs (compare F´´ and G´´ with E´´ or red nuclear signaling in corresponding insets); **H) and I)** Quantification of ISCs/EBs and EEs in control, mad2RNAi and mps1RNAi flies at different timepoints; Tukey boxplot, **** p<0.0001, ** p<0.01, * p<0.05, Mann–Whitney U test; N=28 (intestines) for controls 10d, N=22 for mad2RNAi 10d, N=25 for mps1RNAi 10d, N=22 for mps1RNAi 12d, N=21 (intestines) for controls 20d, N=26 for mad2RNAi 20d, N=21 for mps1RNAi 20d; 3-4 images per intestine used for quantifications; **J) to L)** same genotypes/timepoints as in E) to G) but stained for pH3 to label mitotic cells (white circles or yellow arrows); **M)** quantification of mitotic cells present in the first two fields of view (40X objective in situations) in J-L, mean +/- St Dev; **** p<0.0001, Mann–Whitney U test; N=39 flies for controls, N=24 flies for mad2RNAi, N=17 flies for mps1RNAi; scale bar on A) = 40μm; with the exception of insets, all images are on the same magnification.

The experiments described above (Figure 1) were carried out using *mad2* or *mps1* mutant flies, and, therefore, it is possible that the adult intestinal phenotype we observed could be due to earlier developmental defects. To directly characterize the impact of loss of SAC proteins in ISCs/EBs, we used the esgGAL4 driver to express UAS-RNAi constructs against *mad2* or *mps1* genes. In order to avoid developmental defects, a *GAL80*^*ts*^ (temperature sensitive repressor of the GAL4-UAS system) was used and crosses were maintained at 18°C to suppress the GAL4-UAS system and RNAi mediated knockdown until eclosion when adult flies were shifted to 29°C. In order to test if RNAi mediated knockdown of *mad2* or *mps1* efficiently compromised the SAC in ISCs, flies were fed colchicine as described before and the total number of mitotic cells per ISCs/EBs was determined (Figure S2 A-F, and K). As expected, control flies showed a dramatic increase in pH3 positive cells after colchicine feeding while mad2RNAi or mps1RNAi flies showed a severely compromised mitotic arrest.

To investigate the consequences of a continued SAC impairment, different cell types were quantified in control intestines and intestines where Mad2 or Mps1 proteins were depleted in ISCs/EBs cells by RNAi expression. A significantly higher number of ISCs/EBs was observed in Mad2 or Mps1 depleted cells after 10 days or 12 days of the respective RNAi expression. This accumulation of ISCs/EBs was also observed in 20 day-old mad2RNAi and mps1RNAi flies (Figure 2 E-H). In addition to higher numbers of ISCs/EBs, an accumulation of cells of the EE lineage was also observed (Figure 2 E-G and I), similarly to other models in this tissue where dysplasia was observed [34]. The orientation of cell division of the ISCs has been shown to be subject of biological regulation and it has been shown to impact the stem cell fate decision [13][17][35]. Since an accumulation of ISCs/EBs and EEs has been described previously in a Notch mutant phenotype, and this phenotype was proposed to be mediated by alterations in the orientation of stem cell division [17], we investigated if stem cell division angle in mad2RNAi and mps1RNAi flies was affected. After extensive 3D reconstruction of dividing ISCs, no significant differences were found in the angle of division after Mad2 or Mps1 knockdown when compared to control ISCs (Figure S3 A-H). This clearly excludes this indirect effect as the cause for the dysplasia observed after aneuploidy induction.

In agreement with the accumulation of ISCs/EBs, we found higher proliferation rates in the intestines of mad2RNAi or mps1RNAi flies (Figure 2 J-M), as well as a higher cell density in the epithelia of these SAC impaired flies (Figure S4 A-D). These allowed us to conclude that SAC impairment via *mad2* or *mps1* loss-of-function results in intestinal dysplasia.

### Depletion of SAC genes in ISCs causes aneuploidy

As mentioned previously, flies where Mad2 or Mps1 were knocked down in ISCs/EBs present an impaired SAC response (Figure S2). Consistent with SAC being impaired, lagging chromosomes were observed after Mad2 or Mps1 depletion, a common feature of SAC compromised mitotic cells and known to be linked to aneuploidy generation (Figure 3 A-C). In order to verify in detail that aneuploidy was induced after SAC impairment in the intestine, we performed multiple methods that allow the identification of specific types of aneuploidy. First, and similar to what was performed for the Mad2 and Mps1 mutants, we conducted FACS analysis, which allows us to score a severe type of aneuploidy (>4n). Flies where either mad2RNAi and mps1RNAi were expressed in the intestinal progenitor cells revealed higher percentage of aneuploid cells when compared to control (Figure 3 D-F). FISH analysis allows the identification of extra-chromosomes within cells and has also been used to detect aneuploidy induction. In *Drosophila* high-levels of chromosome pairing can be present [36], therefore an unambiguous classification of cells as aneuploid with FISH is restricted to cells with more than two FISH signals. Using probes for Chromosomes X and III we detected a higher percentage of aneuploid cells in flies expressing RNAi against either Mad2 or Mps1 (Figure 3 G-M). As both FISH and FACS only allow for the detection of types of aneuploidy that involve chromosome gain, we decided to investigate if chromosome loss might also occur after SAC impairment in ISCs/EBs. To do so, we resorted to an antibody for the fourth chromosome [37], which has previously been used successfully to detect loss of this chromosome after SAC impairment in *Drosophila* developmental tissue [5]. Again, this method revealed that this specific type of aneuploidy was increased in ISCs/EBs of intestines of flies where RNAi targeting Mad2 or Mps1 were expressed in progenitor cells (Figure 3 N-Q). Our data indicates that specific knockdown of Mad2 and Mps1 in ISCs/EBs is consistent with the data we obtained with homozygous mutants for these two SAC proteins. These results demonstrate that SAC impairment in intestinal progenitor cells results in aneuploidy induction.

**Figure 3.**
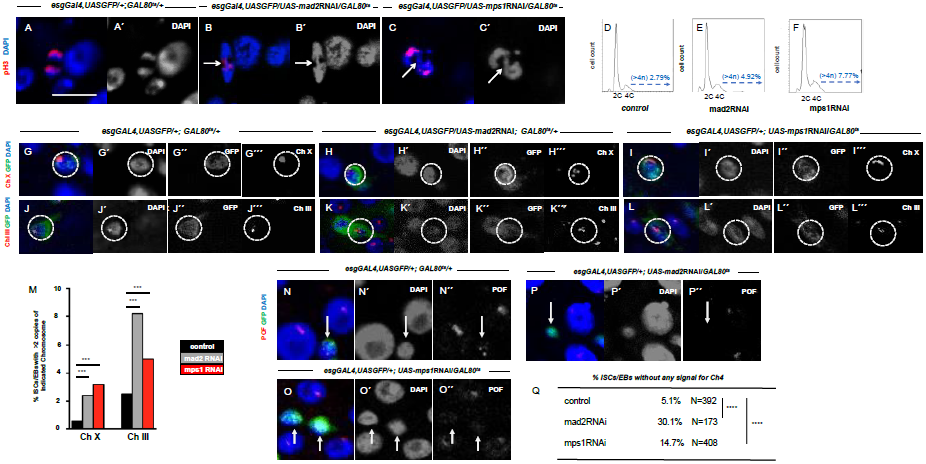
Aneuploidy is generated after SAC impairment. **A)** Control cell in anaphase; **B) and C)** Lagging chromosomes (white arrows) in dividing cells from 10 day-old mad2RNAi and mps1RNAi flies; scale bar in A=2 μm; **D) to F)** Examples of FACS profiles of GFP positive cells (ISCs/EBs) from control, mad2RNAi or mps1RNAi flies; for control and mad2RNAi 2 biological replicates were performed, in both the percentage of aneuploidy was higher in the mad2RNAi flies (p-value<0.01, Fisher exact t-test); for control and mps1RNAi flies 2 biological replicates were performed, in both the percentage of aneuploidy was higher in the mps1RNAi (p-value<0.01, Fisher exact t-test); average of % aneuploidy +/- st dev: control 2.4 +/- 0.5, mad2RNAi 5.4 +/- 0.73, mps1RNAi 7.77 +/- 1.1; **G) to L)** FISH analysis in combination with immunofluorescence allowed us to label chromosomes X or III within ISCs/EBs (GFP+/esg+; white circles); due to somatic chromosome pairing, cells were only scored as aneuploid when more than 2 FISH signals were observed (see examples for mad2RNAi and mps1RNAi in H, I, K, L); **M)** quantification of ISCs/EBs where more than two FISH signals for Chromosome X or III were detected within control intestines and intestines where Mad2 or Mps1 were depleted in ISCs/EBs; ***p<0.001 Fisher exact t-test. **N) to O)** 15d-20d control, mad2RNAi and mps1RNAi intestines stained for anti-POF antibody to label the fourth chromosome; **Q)** percentage of ISCs/EBs where no signal for anti-POF could be observed; N refers to number the ISCs/EBs analyzed in each genotype.

### Aneuploidy causes accumulation of ISCs

Thus far, we have shown that downregulation of SAC genes in ISCs and EBs leads to aneuploidy and a consequent accumulation of these progenitor cell types. Since the esgGAL4 driver is expressed in both ISCs and EBs, we considered important to determine the specific contribution of stem cells to this phenotype. With this purpose, we used the Su(H)LacZ reporter a common marker of EBs [38], and quantified the two different cells types after RNAi induction (Figure 4 A-E). This strategy allowed us to separate these two cell populations and revealed that the percentage of ISCs increased significantly when SAC genes were impaired. Accordingly, we conclude that a deficient SAC response leads to the accumulation of ISCs.

**Figure 4.**
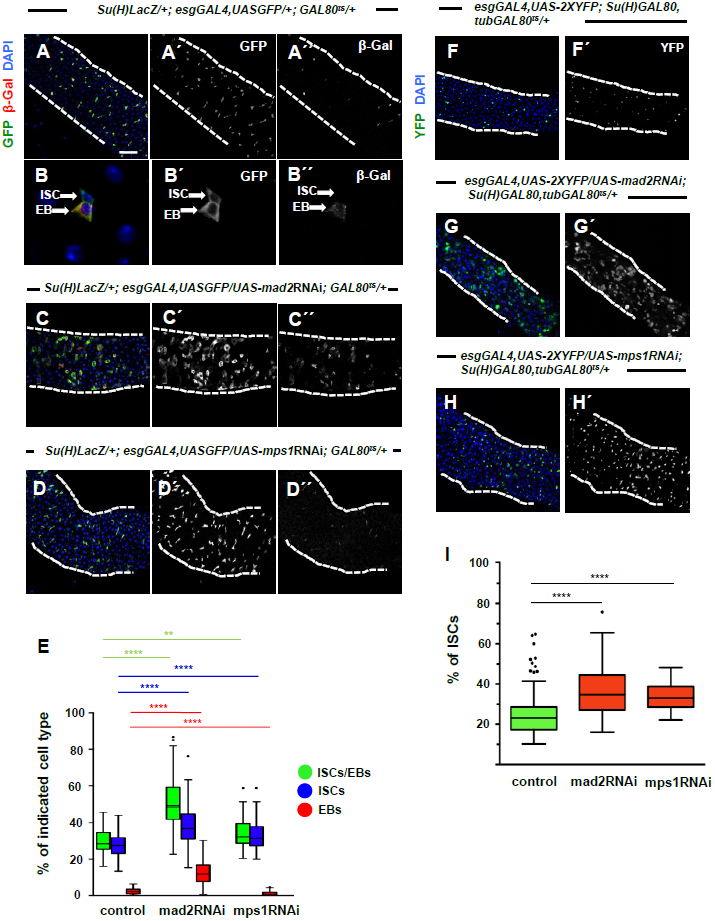
ISCs accumulate after mad2 or mps1 RNAi mediated knockdown. **A)** low and **B)** high magnification of a 20 day-old control intestine where ISCs and EBs can be distinguished based on expression of the Su(H)LacZ protein; *esg* marks ISCs and EBs and beta-Gal marks EBs; **C) and D)** Intestines where RNAi constructs against *mad2* or *mps1* were expressed under the control of the *esgGAL4* promoter during the first 20 days of the *Drosophila* adult and where ISCs and EBs can be distinguished as described for A) and B); **E)** Quantification of percentage of ISCs/EBs (GFP positive, green), ISCs (GFP positive/beta-Gal negative, blue), and EBs (GFP positive/beta-Gal positive, red), per total cell number (DAPI) in control and SAC loss-of-function situations; N=25 for controls, N=25 for mad2RNAi, N=21 for mps1RNAi (intestines); 3-4 images per intestine were used for quantifications; **F)** Control intestine where ISCs can be identified by expression of YFP; **G) and H)** intestines where RNAi against the genes *mad2* or *mps1* were expressed specifically in ISCs during the first 20 days post eclosion; **I)** Quantification of ISCs in situations from F) to H); Tukey boxplot **** p<0.0001, Mann– Whitney U test; N(intestines)=19 for control, N(intestines)=23 for mad2RNAi, and N(intestines)=8 for mps1RNAi; 3-4 images per intestine were used for quantifications; scale bar in A)= 40μm, with the exception of B) all other panels are on the same magnification.

In order to further validate these results, we decided to perform an experiment in which only ISCs would express the RNAi constructs. This was accomplished by using an esgGAL4 driver in combination with a GAL8O^ts^ (expressed in all cells, and temperature sensitive) and a Su(H)GAL80 (expressed in EBs only, and not temperature sensitive), as described in [39]. This strategy allowed us to block expression of the GAL4-UAS system in all cells during development by keeping the flies at 18°C, and activate expression of these constructs, specifically in ISCs, after eclosion, by shifting flies to 29°C. As expected, an accumulation of SAC impaired ISCs was found (Figure 4 F-I), similarly to what we described both for homozygous mutant flies or RNAi expression in ISCs and EBs.

### BubR1 but not Bub3 loss-of-function recapitulate the dysplastic phenotype observed after SAC impairment

Our results described so far contrast with a previous study in which it was proposed that aneuploidy leads to ISCs loss caused by premature differentiation [33]. However, in that study, aneuploidy was induced by RNAi depletion of a different SAC gene (*bub3*). In order to better understand the impact of aneuploidy in ISCs, we analyzed the tissue consequences of loss-of-function of another SAC gene (*bubr1*), and also re-evaluated the *bub3* loss-of-function phenotype in ISCs. An efficient SAC impairment was observed in bubR1RNAi and bub3RNAi flies (Figure S2 G-K). Our results show that bubr1RNAi phenocopies mad2RNAi or mps1RNAi, showing accumulation of ISCs/EBs and of cells of the EE lineage (Figure 5 A-B, E-F). However, bub3RNAi flies show a different phenotype, displaying a reduced number of ISCs/EBs (Figure 5 C, E-F). To exclude the possibility that the bub3RNAi phenotype was caused by an off-target effect of the RNAi construct, we tested a different RNAi line (bub3RNAi^2^) obtaining similar results (Figure 5 D, E-F). Therefore, we were able to reproduce the previously reported detrimental effect of *bub3* depletion in ISCs maintenance [33]. The different phenotype observed with bub3RNAi might be explained by a SAC-independent role of Bub3, which has been shown to form a complex with histone deacetylases, and this interaction appears to confer transcriptional repressor activity during interphase [40]. It is possible that within the genes transcriptionally repressed by this complex some might affect stem cell maintenance.

**Figure 5.**
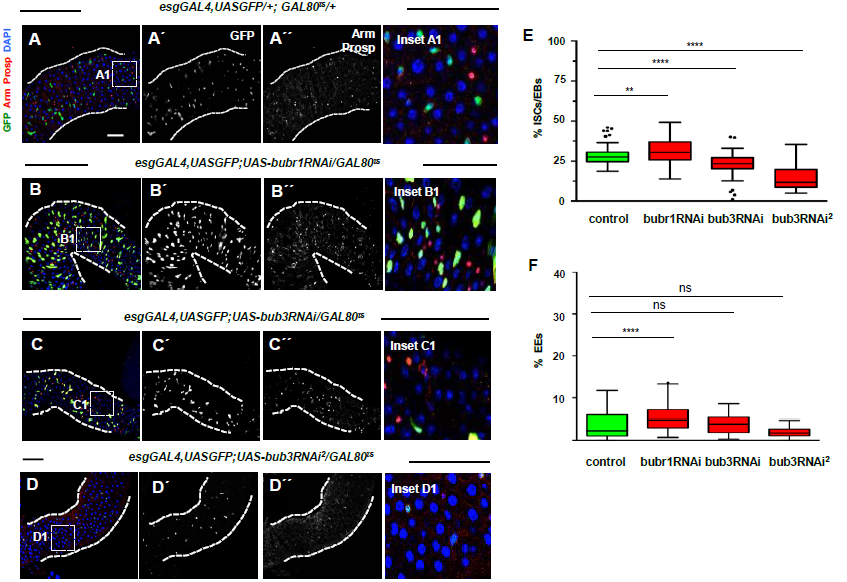
Dysplastic phenotype observed after *mad2* or *mps1* knockdown is recapitulated after *bubr1* loss-of-function but not after *bub3* knockdown. **A)** 20 day-old control intestine where ISCs and EBs are marked based on *esg* expression (green, panel A´), EEs can be identified by expression of prospero (nuclear red signal, A´´); armadillo marks all cells membrane, DAPI marks cell nuclei; **B) to D)** Intestines where RNAi constructs against SAC genes *bubr1* or *bub3* were expressed under the control of the *esgGal4* promoter during the first 20 days of the *Drosophila* adult; note accumulation of ISCs/EBs after BubR1 knockdown (compare B´ with A´) and ISC/EB loss after Bub3 knockdown (compare C´ or D´ with A´); in all genotypes suppression of the GAL4-UAS system was performed during development by using the temperature sensitive repressor GAL80^ts^ and by raising the flies at 18°C; **E) and F)** quantification of ISCs/EBs and EEs from conditions A) to D); Tukey boxplot **** p<0.0001, ** p<0.001, Mann–Whitney U test; N(intestines)=21 for controls, N(intestines)=24 for bubr1RNAi, N(intestines)=19 for bub3RNAi, N=10 for bub3RNAi^2^; 3-4 images per intestine were used for quantifications; scale bar in A) = 40μm, all images are on the same magnification, with the exception of insets.

Interesting parallels can be drawn between our findings and other studies on mammalian models. Consistent with a model where SAC deficiency and aneuploidy induction potentiate tumor development, *mad1* and *mad2* heterozygous offspring showed increased incidence of spontaneous tumors [41-42]. However, haploinsufficiency of *bub3* seems to contrast with other SAC genes loss-of-function phenotypes since it does not result in tumorigenesis in mice models [43]. It would be interesting to re-visit these mice models to study how adult stem cells, in different organs, are affected. Moreover, if available, tools to impair the SAC and induce aneuploidy specifically in stem cells, in a cell-type and temporally-controlled manner, should be explored in these mammalian tissues.

### Apoptosis block delays but does not prevent intestinal dysplasia after aneuploidy induction

In the posterior midgut, ISCs have been shown to be highly sensitive to signals secreted by neighboring cells undergoing cell death or damage, and promote an increased proliferation rate as part of dynamic regenerative process [44]. Therefore, we addressed the possibility that induction of aneuploidy in stem cells could result in apoptosis induction in progenitor cells or in differentiated progenitor cells, and this could result in the activation of a regenerative response of the tissue. We first investigated if apoptotic cells were detected after aneuploidy induction. Pyknotic nuclei were not detected after aneuploidy induction, and the majority of control or mad2RNAi intestines had no cells positive for cleaved caspase-3 (Figure S5 A-C). Bleomycin, a drug known to induce apoptosis in the midgut [45], was used for a positive control for the cleaved caspase-3 antibody.

To better address a putative role for apoptosis in the dysplastic phenotype observed after aneuploidy induction, we genetically impaired the apoptosis response. First, we blocked apoptosis specifically in ISCs/EBs, expressing the baculovirus protein P35. At 10 days, dysplasia was less pronounced when P35 was co-expressed mad2RNAi when compared to mad2RNAi alone, suggesting that blocking apoptosis in ISCs/EBs does not prevent the accumulation of ISCs/EBs but it contributes partially for this phenotype at earlier timepoints (Figure 6 A-C, F). At 20 days, co-expression of P35 with mad2RNAi or mps1RNAi did not prevent the accumulation of ISCs/EBs, and in the case of mps1RNAi, the accumulation of ISCs/EBs was even higher (Figure 6 A-C, F). Interestingly, both at 10 days and 20 days, blocking apoptosis in ISCs/EBs resulted in a more pronounced accumulation of EEs (Figure 6 G). We also impaired apoptosis in all intestinal cell types, by inducing aneuploidy in a background where flies are heterozygous for a deletion in chromosomal deletion Df(3L)H99 that lacks *reaper, head involution defective* and *grim*, three apoptotic inducers. Heterozygous flies for this deletion have been shown to have severely compromised apoptotic response and rescue apoptosis mediated phenotypes [46-50]. At 10 days, expression of mad2RNAi in flies heterozygous for Df(3L)H99 did not result in an accumulation of ISCs/EBs, but a strong accumulation was detected at 20 days in both mad2RNAi or mps1RNAi conditions (Figure 6 A, D-G). A general trend for a stronger accumulation of EEs was also observed in this H99Df(3L) heterozygous background flies for both mad2RNAi and mps1RNAi conditions (Figure 6, D-G). Therefore, the results taken together indicate that blocking apoptosis block does not prevent the dysplastic phenotype, namely the accumulation of ISCs/EBs and EEs over time.

**Figure 6.**
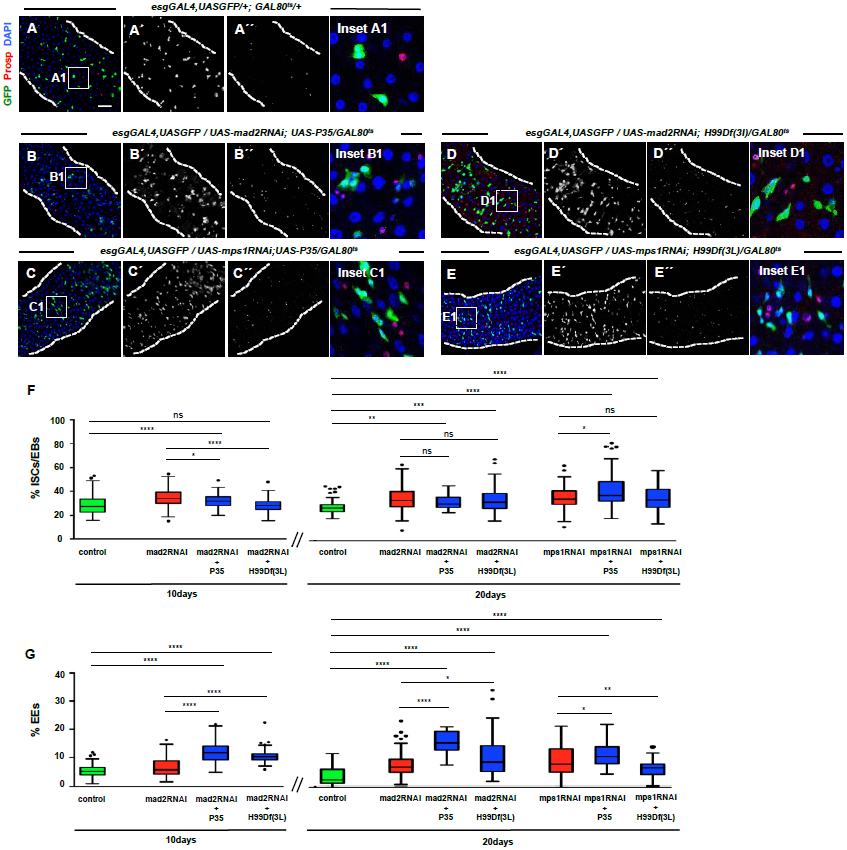
Apoptosis block does not prevent dysplasia observed after aneuploidy induction. **A) to E)** 20 day-old intestines from controls or flies where mad2RNAi or mps1RNAi constructs were expressed in ISCs/EBs and apoptosis was blocked as indicated in respective genotypes; **F) and G)** Quantification of ISCs/EBs or EEs in conditions described in A) to E); control data is the same as shown in Figure 2; control, mad2RNAi and mps1RNAi data are the same as shown in Figure 2; N=20 for 10d mad2RNAi+P35, N=22 for 10d mad2RNAi+H99Df(3L), N=10 for 20d mad2RNAi+P35, N=16 for 20d mad2RNAi+H99Df(3L), N=12 for 20d mps1RNAi+P35, N=16 for 20d mps1RNAi+H99Df(3L); Tukey boxplot **** p<0.0001, *** p<0.001, ** p<0.01, * p<0.05, Mann–Whitney U test; 3-4 images per intestine used for quantifications; scale bar on A)= 40μm; with the exception of insets, all images are on the same magnification.

### JNK autonomous activation in ISCs/EBs modulates intestinal tissue response to aneuploidy induction

Chromosome segregation errors have been shown to induce a DNA damage response within cells [51-52]. Epithelial damage has been shown to cause JNK activation in the *Drosophila* intestine [34][53-55], a pathway that has been shown to mediate the development of a tumorigenic phenotype after aneuploidy induction in *Drosophila* imaginal discs [5]. Based on this, JNK represents a strong candidate to be implicated in the development of intestinal dysplasia after aneuploidy induction. To determine the level of JNK activation we used the reporter line puckered-LacZ [56]. In control intestines, we observed a very low signal in very few cells per intestine (Figure 7 A-B), in sharp contrast to intestines where aneuploidy was induced, where the majority of cells across the entire intestinal section showed strong activation of JNK in both progenitor cells and differentiated cells (Figure 7 C-F). Expression of a dominant negative form of the protein Basket (Bsk^DN^) within ISCs/EBs, is sufficient to prevent JNK activation in these cells and rescue dysplastic phenotypes associated with cellular stress [54][55]. Importantly, expression of Bsk^DN^ *per se* does not impact epithelia homeostasis and stem cell behavior. To test if JNK autonomous inactivation within ISCs/EBs could rescue the dysplastic phenotype, Bsk^DN^ was co-expressed together with mad2RNAi (Figure 7). At 10 days, downregulation of JNK in ISCs/EBs lead to a partial rescue, while at 20 days we could observe a full rescue of the progenitor cell accumulation (Figure 7 G-H). This allowed us to conclude that JNK upregulation within ISCs/EBs is required for the development of intestinal dysplasia after aneuploidy induction. Although our data is strongly supportive of a model where aneuploidy drives intestinal dysplasia in an autonomous fashion within ISCs/EBs, future studies could focus in a putative impact on intestinal homeostasis of the generation of aneuploid genotypes within the differentiated cells of this tissue. Furthermore, as we observe JNK activation in all cells, progenitor and differentiated, it would be interesting to investigate how this general tissue response is regulated and if specific cytokines are produced differently by the different cell types.

**Figure 7.**
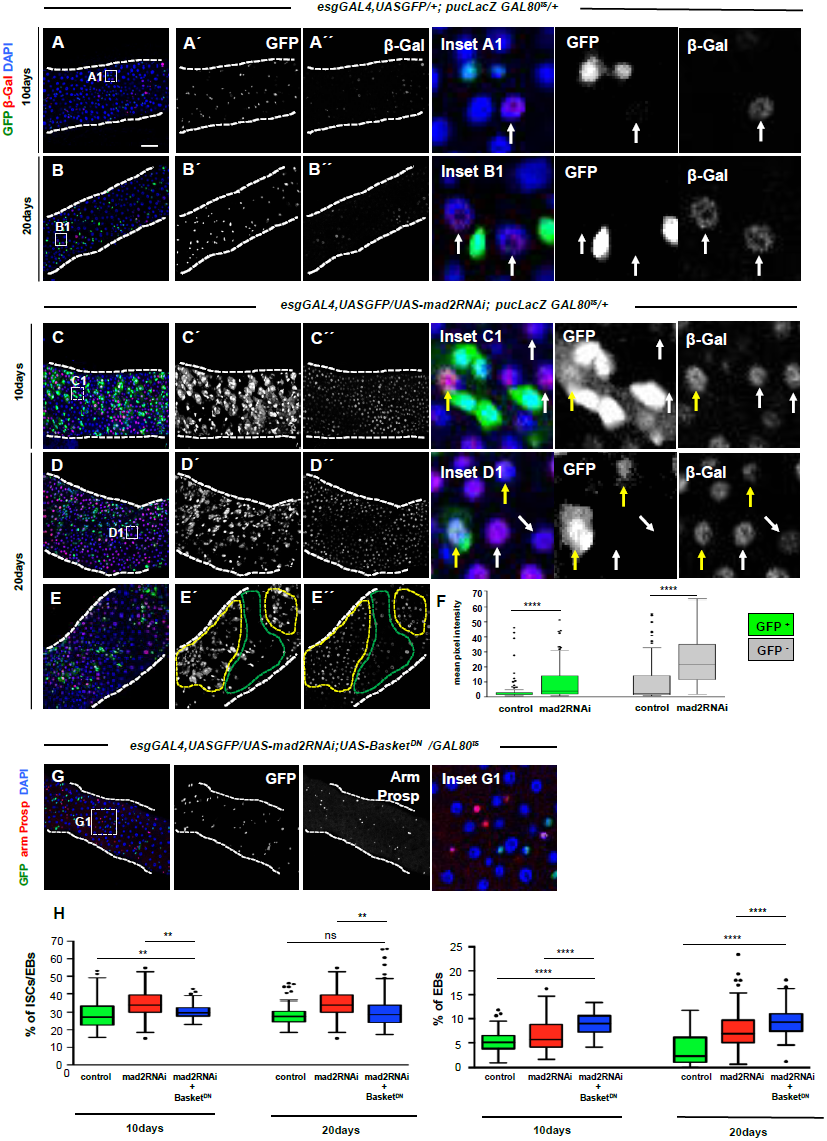
JNK pathway is up-regulated after aneuploidy induction and prevention of this up-regulation within ISCs/EBs partially rescues intestinal dysplasia. **A) and B)** 10 and 20 day-old control intestines where *pucLacZ* serves as readout of the activation status of JNK pathway; *pucLacZ* positive cells were found in low intensity and in very reduced numbers of cells intestine (see A´´ and B´´; white arrows in the insets show examples of *pucLacZ* positive cells; **C) and D)** 10 and 20 day-old intestines where mad2RNAi was expressed and where a strong upregulation of *pucLacZ* can be noted (Compare C´´and D´´with A´´and B´´); up-regulation was found in both progenitor cells (escargot positive, yellow arrows in the insets) and differentiated cells (escargot negative, white arrows in the insets); **E)** 20 day-old intestine where mad2RNAi was expressed; note that *pucLacZ* is upregulated within midgut regions where dysplasia is more pronounced (yellow dashed areas) as oppose to regions within the same gut where dysplasia is absent or less pronounced (green dashed area); **F)** quantification of *pucLacZ* signal intensity in ISCs/EBs (esg/GFP positive) and differentiated cells (esg/GFP negative) at 20 days; scale bar on A) = 40μm; with the exception of insets, all images are on the same magnification; **G)** 20 day-old intestine where mad2RNAi was co-expressed with Bsk^DN^ in ISCs/EBs; note absence of dysplasia; compare with control and mad2RNAi in Figure 2; **H)** quantification of the percentage ISCs/EBs and EEs in control, mad2RNAi and mad2RNAi + Basket^DN^; control and mad2RNAi data are the same as shown in Figure 2; N=14 for 10d mad2RNAi+basket^DN^, N=21 for 20d mad2RNAi+basket^DN^; 3-4 images per intestine used for quantifications; Tukey boxplot **** p<0.0001, ** p<0.01.

Interestingly, the accumulation of EEs was not rescued neither at 10d or 20d when Bsk^DN^ was co-expressed with mad2RNAi, suggesting that accumulation of EEs after aneuploidy induction is independent of progenitor cell accumulation (Figure 7 G,H). Further studies should address which biological mechanisms are responsible for the accumulation of EEs, after aneuploidy induction.

### Aneuploidy caused by centrosome or kinetochore impairment results in intestinal dysplasia

In order to confirm that intestinal dysplasia is caused by aneuploidy and not SAC impairment *per se*, we induced aneuploidy in ISCs either by impairing kinetochore function (and, consequently, microtubule attachment) or by inducing centrosome amplification. Successful induction of aneuploidy in *Drosophila* has been previously achieved by knockdown of the centromeric-associated protein meta (Cenp-meta), a kinesin-like motor protein required for efficient end-on attachment of kinetochores to the spindle microtubules [57-58]. Cenp-E knockout mice (the mammalian homolog of Cenp-meta) have also been shown to accumulate aneuploid cells [59]. Thus, we depleted Cenp-meta by expressing an RNAi construct in ISCs/EBs using the esgGAL4 driver. After 20-days of RNAi expression, we observed an accumulation of ISCs/EBs and cells of the EE lineage, phenotype in all respects similar to the one observed after loss of SAC genes (Figure 8 A-B, D-E). To analyze this further, we induced alterations in cell division through centrosome amplification since these have been associated with aneuploidy and tumor development in both *Drosophila* and humans [60-61]. In *Drosophila*, overexpression of the Sak kinase (Sak), the *Drosophila* homolog of PLK4, leads to centrosome amplification and, consequently, aneuploidy [33][62]. Thus, we induced the overexpression of Sak in ISCs/EBs using the esgGAL4 driver. The results showed a strong accumulation of ISCs/EBs and EEs, similarly to what was observed for loss of SAC genes or depletion of Cenp-meta (Figure 8 A, C-E). Higher proliferation rates and specific ISCs accumulation was found in intestines where these Cenp-meta was depleted or Sak was over-expressed (Figure 8 F-I and Figure S6 A-D), similarly to what we observed when aneuploidy was induced via SAC impairment. Thus, we conclude that the dysplastic phenotype we observed in the midgut can be caused by impairment of different biological mechanisms/structures that result in the induction of aneuploidy (Figure 8 J).

**Figure 8.**
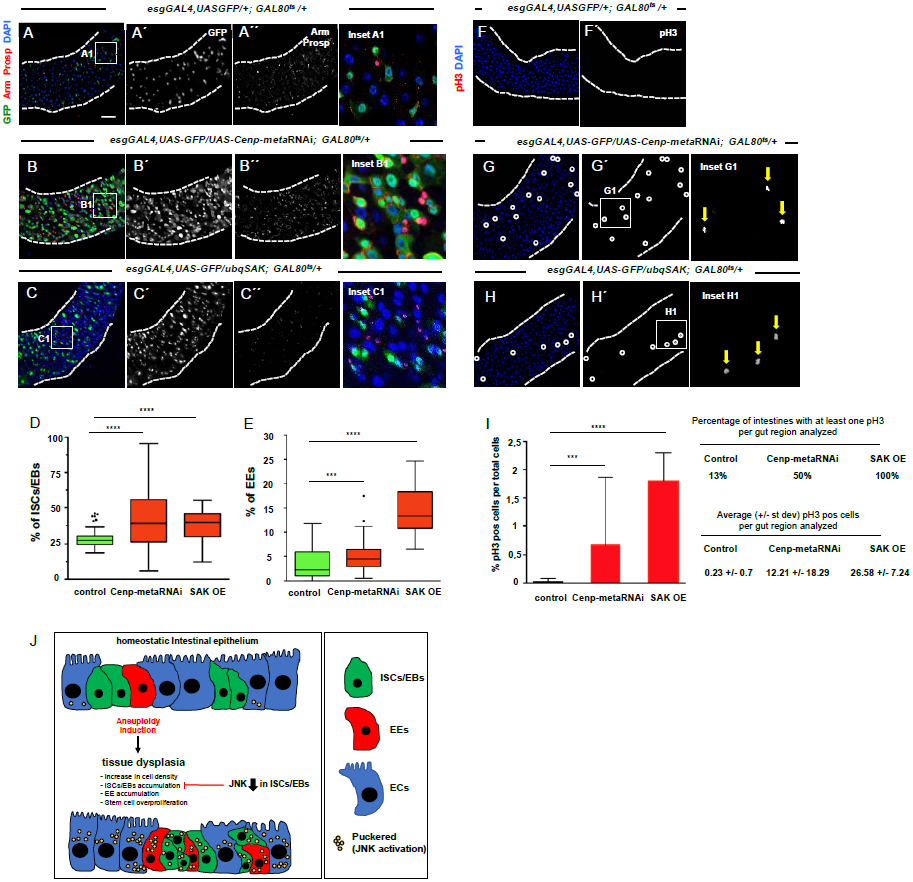
Aneuploidy induction in ISCs via impairment of kinetochore-microtubule attachment or centrosome amplification leads to intestinal dysplasia. **A)** 20 day-old control intestine where ISCs and EBs are marked based on *esg* expression (green, panel A´), EEs can be identified by expression of prospero (nuclear red signal, A´´), armadillo (Arm) marks all cells membrane, DAPI marks cell nuclei; **B)** Example of an intestine where RNAi construct against *cenp-meta* was expressed under the control of the esgGAL4 driver during the first 20 days of the *Drosophila* adult; **C)** Example of a 20 day-old intestine where the protein Sak was constitutively overexpressed (OE); **D and E)** Quantification of ISCs/EBs and EEs cells in situations from A) to C); control data is the same as shown in Figure 2; Tukey boxplot **** p<0.0001, Mann–Whitney U test; N(intestines)=21 flies for control, N(intestines)=32 flies for cenp-metaRNAi, and N(intestines)=8 for Sak overexpression; 3-4 images per intestine used for quantifications; **F) to H)** same genetic conditions and time point as described for A) to C) but stained for pH3; **I)** quantification of mitotic cells present in the first two fields of view (40X objective in situations) in F-H, mean +/- St Dev; **** p<0.0001, *** p<0.001, Mann–Whitney U test; N(intestines)=39 flies for control (control data is the same as in Figure 2), N(intestines)=18 flies for cenp-metaRNAi, and N(intestines)=24 for Sak overexpression; scale bar on A) = 40μm; with the exception of insets, all images are on the same magnification; **J)** Graphical abstract of results gathered on the impact of aneuploidy induction in the intestine epithelium.

We describe here *in vivo* examples of how stem cell failure to maintain genomic stability can lead to tissue pathology. Current knowledge strongly implies that the effect of aneuploidy on adult stem behavior is tissue specific, therefore a cautioned extrapolation of our findings to other *Drosophila* or mammalian adult tissues should be performed. Nevertheless, we highlight one specific tissue example where mechanisms that ensure correct chromosome segregation during stem cell proliferation prevent tissue dysplasia. Future studies should focus on whether *Drosophila* and mammalian adult stem cells don´t activate cell-death mechanisms in response to aneuploidy, on *Drosophila* ISCs mechanisms of resistance to aneuploidy, and on the identification of the specific genomic alterations that drive accumulation of progenitor cells. Moreover, the depth of our experiments does not allow us to exclude the possibility that some specific types of aneuploidy might have a detrimental effect and even cause ISC death, so it will be important to determine which specific types of aneuploidy are tolerated by stem cells. Mammalian and *Drosophila* intestinal epithelia share many common similarities that include cell type composition, anatomic compartmentalization, and pathways governing tissue maintenance and regeneration [63]. Furthermore, *Drosophila* has given significant contributions to our understanding of the development of different human pathologies,including diseases that are characterized by an accumulation of progenitor cells and higher proliferation rates like cancer [64]. Therefore, it would be important to determine whether mammalian adult gut displays similar responses to aneuploidy induction within stem cells.

## Author Contributions

L.P.R. and C.E.S planned experiments. L.P.R., A.M., R.B., and T.L. performed the experiments and together with C.E.S performed data analysis. L.P.R. and C.E.S wrote and edited the manuscript.

## Acknowledgements

The authors thank Leanne Jones, Carla Lopes, Bruce Edgar, Paulo Pereira, Jan Larsson, and S. Hou laboratories, and the Vienna Drosophila RNAi Center for reagents and stocks. This article is a result of the project Norte-01-0145-FEDER-000029 - Advancing Cancer Research: From basic knowledge to application, supported by Norte Portugal Regional Operational Programme (NORTE 2020), under the PORTUGAL 2020 Partnership Agreement, through the European Regional Development Fund (ERDF) and it is also funded by National Funds through FCT – Fundação para a Ciência e a Tecnologia undcfer the project PTDC/BEX-BCM/1921/2014

## Abbreviations list

arm: armadillo
cenp-meta: centromeric protein meta
EB: enteroblast
EC: enterocyte
EE: enteroendocrine Esg - escargot
GFP: green fluorescent protein
ISC: intestinal stem cell
pH3: phospho-histone 3 pros - prospero
SAC: spindle-assembly checkpoint
SaK: sak kinase
Su(H): supressor of hairless
YFP: yellow fluorescent protein

## Supplemental Information

Supplemental information includes 6 figures, Supplemental Experimental procedures and Supplemental references.

## Supplementary Information

### Methods

#### Fly husbandry and stocks

Flies were raised on standard cornmeal-molasses-agar medium. Female progeny from experimental crosses were collected and maintained with less than 30 flies per vial. Flies were turned onto fresh food vials every two days. The following fly stocks used were from the Bloomington *Drosophila* Stock Center (BDSC), Vienna Drosophila Stock Center (VDRC), or generous gifts from the fly community as indicated: *mad2*^*Δ*^, *aldG*^*4422*^, *UAS-mad2RNAi, UAS-mps1RNAi (*Bloomington stock center); *esgGal4,UAS-GFP; Gal80*^*ts*^ *and Su(H)LacZ; esgGal4,UASGFP;Gal80*^*ts*^ (gifts from Dr Leanne Jones, UCLA); *esgGal4,2xYFP; Su(H)Gal80, tub-Gal80t*s (gift from Steven Xou, NCI, NIH); pUBGFP-SAK (gift from Dr Renata Basto); *mad2GFP* and *bubr1GFP* (gifts from Dr Roger Karess). *Wild-type* flies were *Oregon R.* More detailed information about these stocks can be found at Flybase (http://flybase.bio.indiana.edu).

#### Immunostaining, microscopy and data analysis

Immunofluorescence (IF) microscopy was performed on whole-mount intestines dissected directly in 4% PFA and left ON at 4°C for fixation. After fixation, three 10-minute washes with PBST (PBS 0,1% triton) were performed and then samples were incubated for 1 hour with a blocking solution of PBST/BSA (PBS 0,1% triton 1% BSA). Then, samples were incubated with primary antibodies ON at 4°C. Then, three 10-minute washes with PBST were performed and intestines were incubated two hours with secondary antibodies protected from light. Lastly, three 10-minute washes with PBST again performed and then intestines were mounted in Vectashield mounting medium with DAPI (Vector Laboratories). Images were obtained using a Leica TCS SP5 II (Leica Microsystems) confocal microscope with HC PL APO CS 40x/NA 1.10 objective and a LAS 2.6 software. All images were acquired on the first 2 fields of view of the posterior midgut (after the pyloric ring) on a 40x water objective - based on recent anatomic and physiological characterizations of the *Drosophila* intestine this corresponds to the P3-P4 regions [S1] or R4-R5 region [S2]. In this work, we adopted sample-sizes that are typically used in the field and for the model system under study. As a general rule, at least 20 intestines were analyzed from at least two biological replicates (progeny from different crosses) and at least 10 of those intestines were used for quantifications. Images were taken from both top and bottom layers of the intestines. The N mentioned in figure legends always corresponds to the number of flies/intestines analyzed, and for each intestine 4 images were taken and used for quantifications: 2 images from the first field of view on the 40x objective (top and bottom) plus two images from the second field of view (top and bottom). Images were analyzed and edited in the following softwares: LAS 2.6, Adobe Photoshop (Mountain View, CA) and ImageJ 1.50i. Statistical analysis and graphical display were performed using the Prism5 (GraphPad) software.

#### Antibodies

Intestines were stained with: mouse anti-armadillo (1:20) and mouse anti-prospero (1:100) (Developmental Studies Hybridoma Bank, developed under the auspices of the National Institute of Child Health and Human Development and maintained by The University of Iowa, Department of Biological Sciences); rabbit anti-phospho-histone H3 (1:2500) (Millipore); rabbit anti-POF (gift by Dr Jan Larsson; 1:500); rabbit anti-GFP (1:5000) (Molecular Probes) mouse anti-β-Galactosidase (Cappel, 55976, MP Biomedicals, 1:1000); rabbit anti-H2AvD (Rockland); rabbit anti-pERK antibody (Phospho-p44/42 MAPK) at 1:200 (4370, Cell Signaling); rabbit anti-cleaved Caspase-3 at 1:200 (9661, Cell Signaling). Secondary antibodies were diluted 1:500 (Molecular Probes)

#### Colchicine treatments

Solutions of either 1) 0,2 mM colchicine with 5% sucrose (experiment), or 2) 5% sucrose only (control) were prepared fresh on each day of experiment and 2000μL of solution were applied to a filter paper that was placed at the bottom of each empty vial. Flies were transferred into these vials with soaked kimwipe after a period of feeding/aging in normal food: 4-6 days post-eclosion in the case of mutants and *wild-type* (Figure 1), and 2 days post-eclosion on the case of the RNAi and controls experiments (Figure S1 and Figure S3). After 24h of feeding in vials with control or colchicine solutions, intestines were fixed and subjected to immunofluorescence in order to evaluate mitotic rates.

#### FACS analysis

Female intestines were dissected (30-50 per genotype) in 1xPBS/1% BSA solution in dissecting slides on ice, for no more than 2h, and transferred to 1.5ml eppendorfs also on ice. After all intestines for different genotypes were dissected, initial sample preparation was performed according to [S3]. Once dissociation of cells was achieved, subsequent protocol steps were then performed as described in [S4]. 3,7% formaldehyde in 1xPBS was used for fixation and propidium iodide (PI) and RNAse incubation was done overnight. PI fluorescence was determined by flow cytometry using a BD FACS Canto II flow cytometer. DNA analysis (ploidy analysis) on single fluorescence histograms was done using FlowJo (TreeStar, Inc.). Experimental conditions were always compared with control samples dissected and FAC sorted in the same day. Multiple biological replicates were always performed, and their number is indicated in the corresponding figure legends.

#### Fluorescence in situ hybridization (FISH)

Oligonucleotides probe for dodeca heterochromatic repeat (Chromosome III) and 359 repeats (Chromosome X) were both synthesized with a 5’CY3, by Integrated DNA Technologies (IDT). The following sequences were used: 5’ CY3-CCCGTACTGGTCCCGTACTGGTCCCG (Chr III) and 5’ CY3-GGGATCGTTAGCACTGGTAATTAGCTGC (Ch X). Our FISH protocol was adapted from previously described methods [S5-S7]. Identification of ISCs/EBs (GFP+) is required. Thus, intestines from female flies were dissected and fixed overnight (O/N) in PFA 4%, washed 3 times in PBS-T (PBS, 0.3% triton), blocked for 30min in PBS 1X/BSA and incubated with an anti-GFP antibody (1:1000) overnight (O/N) washed three times in PBS-T and incubated with secondary antibody 2 hours at room temperature. Intestines were then washed three times in PBS-T and fixed another time for 40min in PFA 4% before proceeding with the probe hybridization. Intestines were then washed three times in PBST, 1X in 2xSSCT (2XSSC (EU0300, Euromedex)/0.1% tween-20), 1X in 2XSSCT/50% formamide (47671, Sigma). For the pre-hybridization step, intestines were transferred to a PCR tube containing 92°C pre-warmed 2XSSCT/50% formamide and denaturated 3min at 92 °C. Intestines were then hybridized 5min at 92 °C with previously denaturated DNA probe (40-80 ng) in hybridization buffer (20% dextran sulfate (D8906, sigma)/2XSSCT/50% deionized formamide (F9037, Sigma), 0.5 mg ml_1 salmon sperm DNA (D1626, Sigma) 3min after denaturation at 92 °C, tubes were left O/N at 37 °C. Samples were then washed with 60°C pre-warmed 2XSSCT for 10 min, and 1×5min in 2XSSCT at room temperature. Probe hybridization in intestinal samples were performed in a thermocycler. Samples were then incubated in Mounting medium for fluorescence with DAPI (H-1200 from vector Laboratory Inc. Burlingame, CA 94010).

In *Drosophila* high-levels of chromosome pairing can be present [S8]. Therefore, for the two chromosome probes used, cells were scored as aneuploid only when more than two dots of the respective FISH probes were detected.

#### Loss of Chromosome IV

Immunostaining with the anti-Painting of Fourth (POF) antibody was used to label the fourth chromosome. ISCs/EBs were identified based on *escargotGal4* expression cells and cells without immunofluorescence signal for the anti-Painting of Fourth antibody were considered as cells that lost Chromosome IV.

#### Determination of mitotic angles

ISCs undergoing mitosis were identified by positive staining of the mitotic marker phospho-histone H3 (pH3). Alfa-tubulin was used in combination with pH3 and DAPI to determine the mitotic phase. Spindle orientation in stem cells is dynamic, and mechanisms regulating its position can act until anaphase is initiated [S9-10]. Therefore, in order to measure the final mitotic angle, cells in prometaphase and metaphase were excluded from the analysis and mitotic angle was only determined in cells at anaphase or telophase. Multiple confocal z-stacks were acquired and three-dimensional reconstructions were performed using ImageJ 1.50i. In these three-dimensional reconstruction images, alfa-tubulin was used to identify the basement membrane. One line was drawn aligned with the basement membrane and another one was drawn intersecting the center of the two separating nuclei of mitotic cell. The angle between these two lines was determined using ImageJ 1.50i, and considered the angle of stem cell division.

## Supplementary Figures

**Figure S1.**
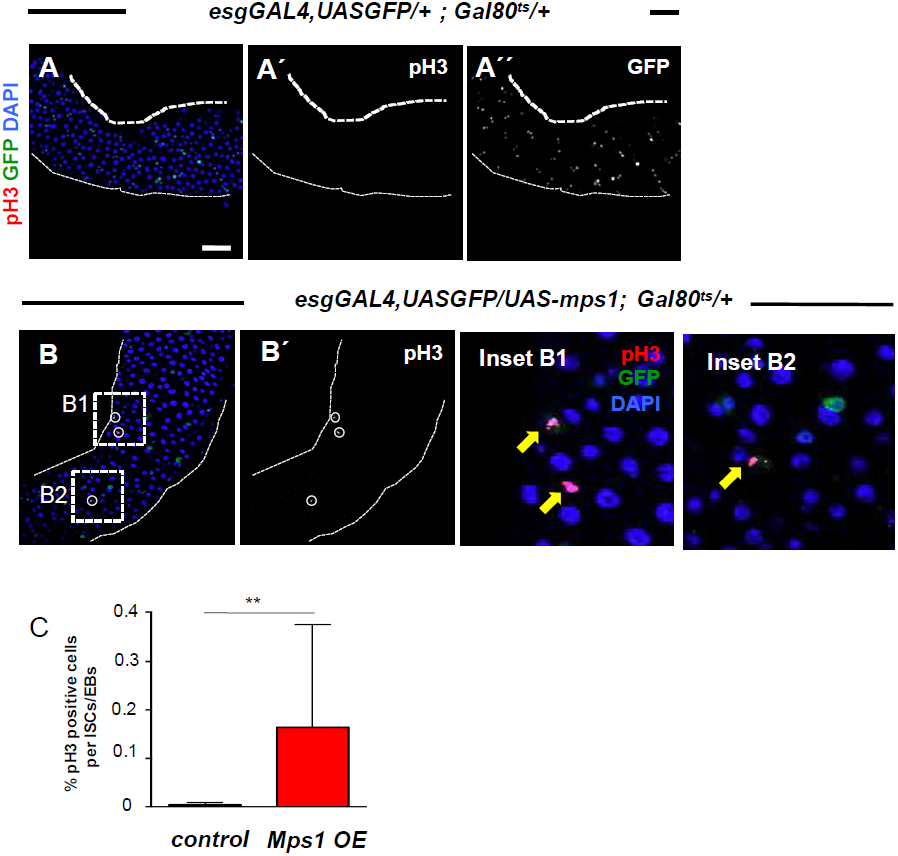
related to Figure 1. Mps1 over-expression leads to ISC cell-cycle arrest. **A)** 2 day-old control intestine stained for GFP marking *esg* positive cells (ISCs/EBs) and pH3 positive cells; in the majority of intestines (20 out of 21 total analyzed) no pH3 were found in the first two-fields of view using a 40X objective; **B)** intestine where Mps1 was over-expressed under the control of the *esgGal4* promoter during the first 2 days post eclosion showing three pH3 positive cells (white circles, insets B1 and B2, yellow arrows); pH3 positive cells were found in the first two-fields of view using a 40X objective in more than half of the intestines analyzed (10 out of 19 total analyzed); **C)** quantification of pH3 positive cells per total cells in situations A) and B); Tukey boxplot, ** p<0.01, Mann–Whitney U test; N(intestines)=21 for controls and N(intestines)=19 for Mps1 over-expression; in both A) and B) GAL4-UAS was suppressed during development using the temperature sensitive repressor GAL80ts and by raising the flies at 18°C.

**Figure S2.**
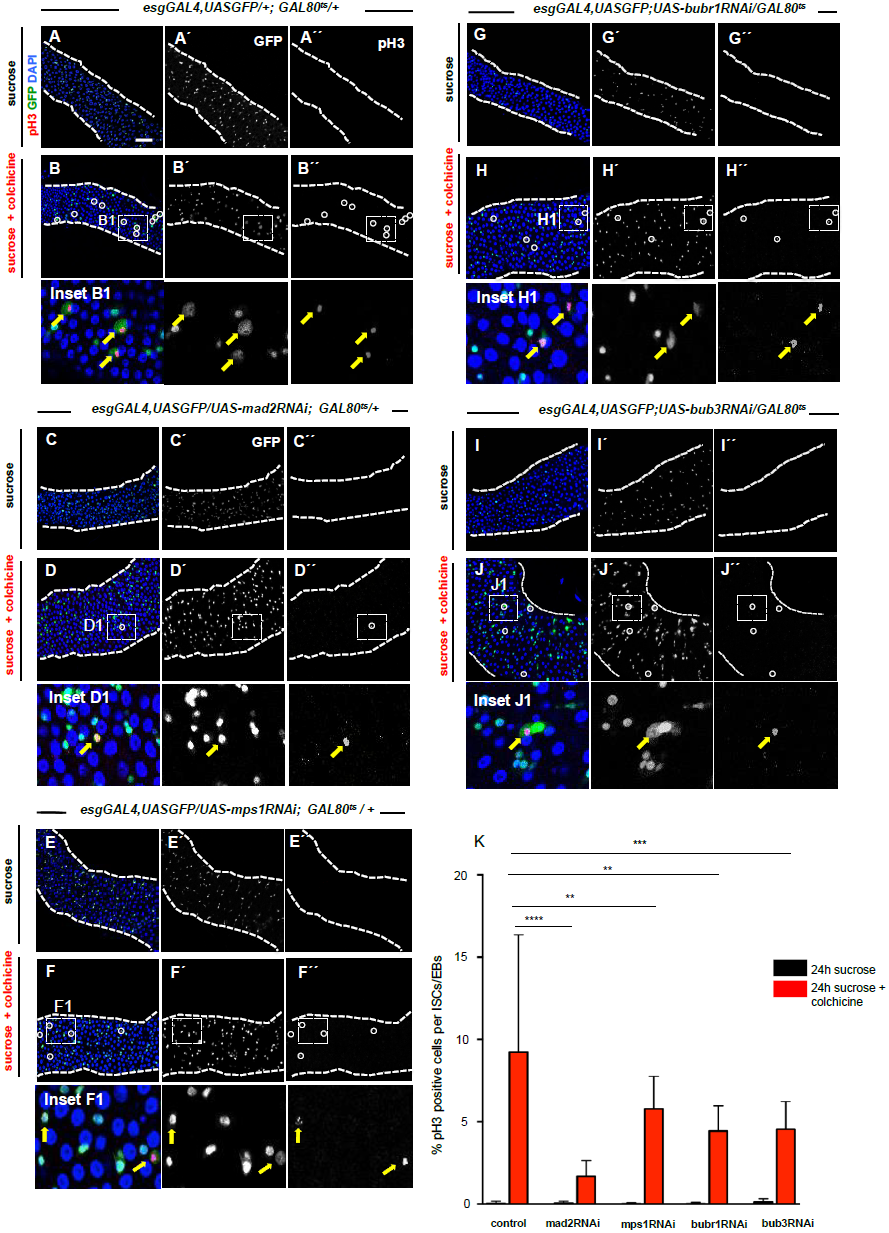
related to Figure 2 and 5. SAC response in ISCs from flies expressing RNAis against Mad2, Mps1, BubR1 or Bub3. **A to J)** Mitotic cells labeled with phospho-histone 3 (pH3, Green) in intestines from control flies and flies where indicated RNAi was expressed; control and RNAi flies were kept at 18°C during development to suppress the GAL4-UAS system and then shifted to 29°C at eclosion day; after 48h at 29°C on regular food, flies were shifted to vials with either sucrose or sucrose+colchicine solutions for 24h; N(intestines)>22 for all genotypes/conditions; scale bar in A)= 40μm, all intestine images are in the same magnification; **K)** % pH3 positive cells per progenitor cells in situations shown from A to J; ** p<0.01, *** p<0.001, **** p<0.0001 Mann–Whitney U test.

**Figure S3.**
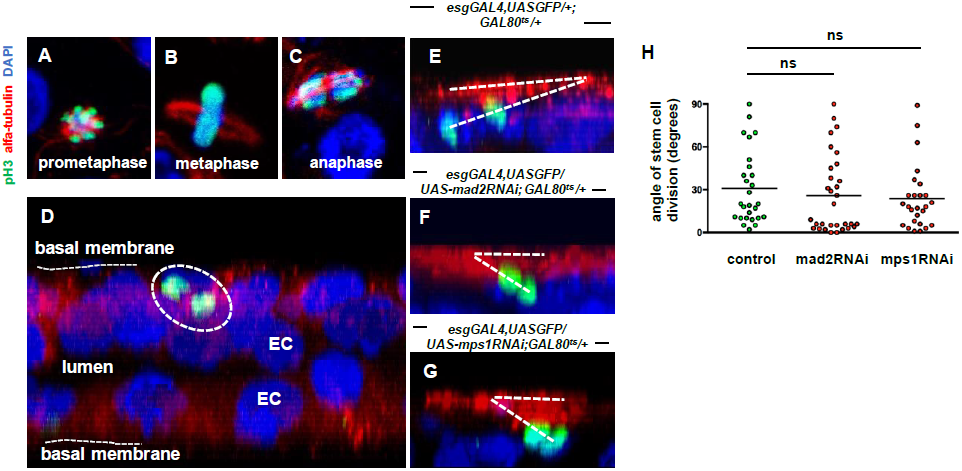
related to Figure 2. SAC impairment does not affect the angle of stem cell division. **A to C)** Different mitotic phases could be identified in 15-20 day-old intestinal samples using pH3 (green) to label mitotic cells, alfa-tubulin (red) to stain the spindle, and DAPI to mark the nuclei (blue); **D)** Example of a broad view of a 3D reconstruction (X axis projection) of multiple z-stacks of an intestinal sample; general architecture of the tissue can be appreciated with the ECs facing the lumen found in between two epithelial layers, and where diving ISC (white ellipse) can be found adjacent to the basement membrane (made evident by alfa-tubulin, red); **E to G)** Examples of diving ISCs in Control, mad2RNAi and mps1RNAi 15-20 day-old intestines; mitotic angle was inferred by the angle formed by two lines drawn: one along the basement membrane and one intersecting the center of the two separating nuclei; **H)** Distribution of mitotic angles determined for three conditions described in E-G; Controls (N=27), mad2RNAi (N=31) and mps1RNAi (N=26); p-value=0.14 controls vs mad2RNAi, p-value=0.29 in controls vs mps1RNAi, Mann–Whitney U test.

**Figure S4.**
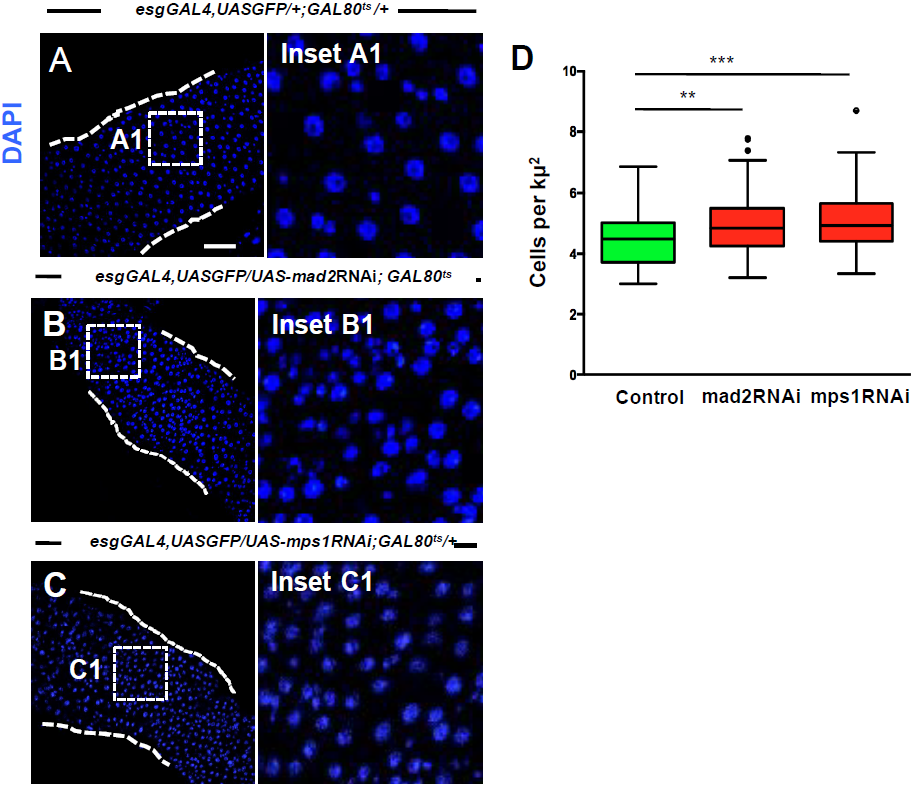
related to Figure 2. Cell density of intestines after SAC impairment by Mad2 or Mps1 knock-down. **A)** 20 day-old control intestine; cell nuclei marked by DAPI; **B) and C)** Intestines where RNAi constructs against SAC genes *mad2* or *mps1* were expressed under the control of the *esgGal4* promoter during the first 20 days of the *Drosophila* adult; note high cell density (compare B1 or C1 with A1); in control and RNAi conditions suppression of the GAL4-UAS system was performed during development by using the temperature sensitive repressor GAL80ts and by raising the flies at 18°C; N(intestines)=20 for controls and N(intestines)=20 for mad2RNAi and N(intestines)=18 for mps1RNAi; **D)** quantification of cell density in situations A to C); scale bar in A)= 40μm, all images are on the same magnification; ** p<0.01, *** p<0.001 Mann–Whitney U test.

**Figure S5.**
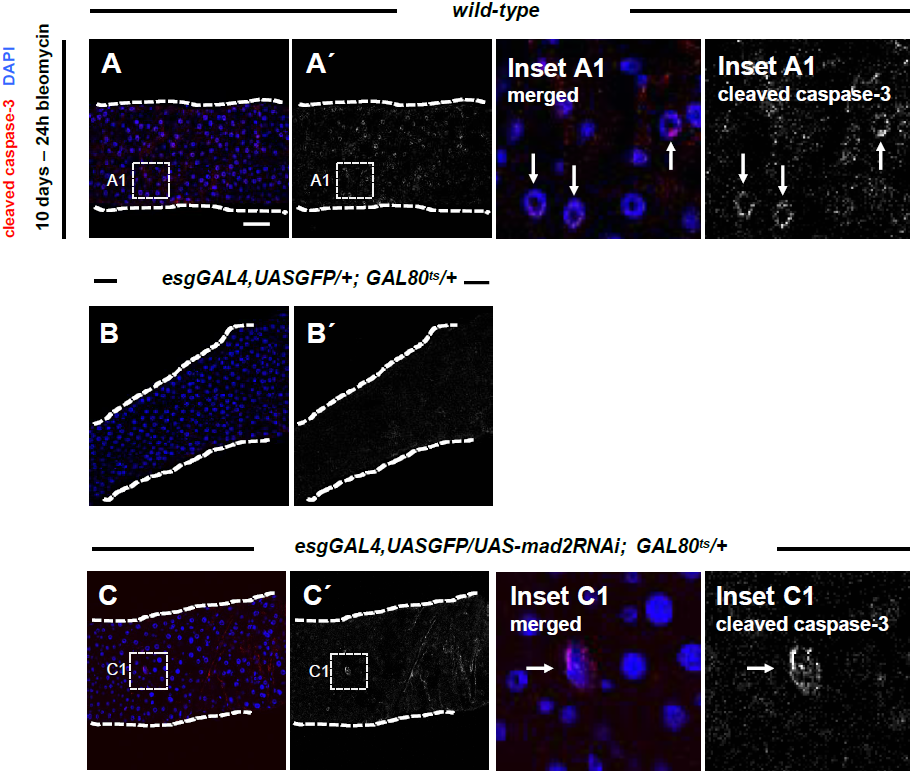
related to Figure 6. Aneuploidy induction does not result in significant activation of apoptosis. **A)** 10 day-old wild-type intestine fed 24h with bleomycin as a positive control for cleaved caspase-3 staining (apoptotic marker); many apoptotic cells were detected (see A´), examples shown in inset and pointed by white arrows; **B) and C)** 10 day-old control and mad2RNAi intestines stained for cleaved caspase-3; no apoptotic cells were found in the control samples, N=10 (intestines); in a total of 17 intestines apoptotic cells were only found in only 2 intestines for mad2RNAi (2 apoptotic cells in one intestine and 3 in the other one); scale bar = 40μm, all images are on the same magnification.

**Figure S6.**
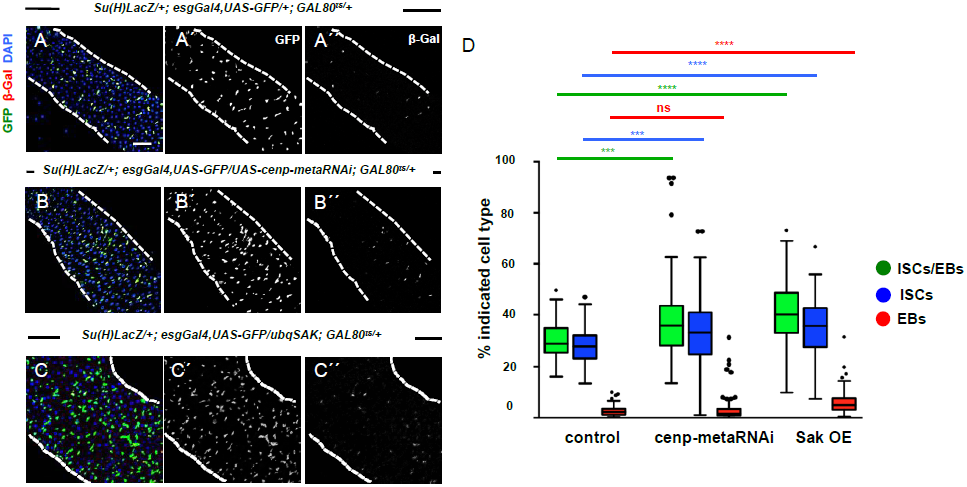
related to Figure 8. ISCs accumulate after Cenp-meta knockdown or Sak overexpression. **A)** 20 day-old control intestine where ISCs and EBs can be distinguished based on expression of the Su(H)LacZ protein like described for Figure 4; **B)** Example of an intestine where a RNAi construct against *cenp-meta* was expressed under the control of the esgGAL4 driver during the first 20 days of the *Drosophila* adult, and **C)** Example of a 20 day-old intestine where the protein Sak was constitutively overexpressed (OE); **D)** quantification of situations A) to C); Tukey boxplot; *** p<0.001, **** p<0.0001, Mann–Whitney U test; N(intestines)>20 for all genotypes; scale bar = 40μm, all images are on the same magnification.

